# A Coupled Flow-Thermoregulation Lumped Model to Investigate Cardiac Function

**DOI:** 10.1101/2021.05.02.442367

**Authors:** Amin Deyranlou, Alistair Revell, Amir Keshmiri

**Affiliations:** Wellcome/EPSRC Centre for Interventional and Surgical Sciences (WEISS), Department of Medical Physics and Biomedical Engineering, University College London, 43-45 Foley Street, London, W1W 7TS, UK; Department of Mechanical, Aerospace and Civil Engineering (MACE), The University of Manchester, Manchester, M13 9PL, UK; Manchester University NHS Foundation Trust, Manchester Academic Health Science Centre, Southmoor Road, Wythenshawe, Manchester, M13 9PL, UK

**Keywords:** Lumped modelling, thermoregulation, cardiovascular flow, numerical method

## Abstract

Lumped (zero-dimensional) technique is a robust and widely used approach to mathematically model and explore bulk behaviour of different physical phenomena in a lower expense. In modelling of cardio/cerebrovascular fluid dynamics, this technique facilitates the assessment of relevant metrics such as flow, pressure, and temperature at different locations over a large network/domain. Furthermore, they can be employed as boundary conditions in multiscale modelling of physiological flows. In this methodology paper, a lumped model for the cardiovascular flow simulation along with a two-node thermoregulation model are employed. The lumped models are built upon previous studies and are amended appropriately to focus on cardiac function. The output of the coupled model can either be used for assessing the cardiac function in different physiological conditions or it can provide the input data for other investigations. Noteworthy to mention that, the present model has been specifically developed for investigation on the effects of atrial fibrillation on cardiac performance.

## Method details

### Background

The lumped modelling technique is a simple, zero-dimensional approach, which enables to mathematically model and solve a wide range of physical phenomena. In modelling of cardiovascular flow, the lumped technique describes general behaviour of a system through distributions of pressure and flow rate of a region of interest. In this method, each compartment of a cardiovascular network is mimicked within a hydraulic impedance, which is the combination of frictional loss, vascular compliance, and blood inertia. There is a wide spectrum of lumped models, which starts from a simple two-element Windkessel, to the most extended one named as Guyton model [1]. Employing lumped techniques for the cardiovascular systems, it brings up several applications such as flow analysis in systemic arteries, haemodynamic responses of a native cardiovascular system under various physiological conditions, and in surgical and therapeutic interventions, ventricular assist device support for the heart failure, study of cardiovascular responses under different neuro-regulation conditions, and as boundary conditions in multiscale modelling. Shi et al. [2] performed a comprehensive review on the lumped techniques in modelling of cardiovascular flow. In a broad classification by Shi et al. [2], they summarised that to generate a circuit of heart and vasculature, several mono and multi-compartment elements are employed, which the latter is again composed of several mono-compartment elements.

Lumped approach is not just limited to cardio/cerebrovascular networks. Examining body thermal response to different physiological and environmental conditions is another application of lumped technique to understand biological performance of human’s body. The thermoregulation model proposed by Gagge et al. [3] is one of the preliminary attempts, which was later on developed by several authors, such as one of the very recent improvement proposed by Zhang et al. [4]. In this model, the body is divided into skin and core regions, hence, it is called two-node thermoregulation model. Then, a set of governing equations for heat and mass transfers, fluid flow, and the body metabolism are devised to estimate internal and skin temperatures of the body in a variety of circumstances.

Building upon our previous workflows in modelling of cardiovascular flow [5]–[13], in this methodology paper, a coupled lumped model for the flow and thermoregulation of the body is employed to evaluate cardiac function in normal and diseased conditions. The following sections are organised to initially present the lumped model for the cardiovascular flow and then a two-node thermoregulation model is described. Finally, the employed methods are validated to evaluate the proposed technique. The developed pipeline will be used for further analysis on one of the cardiovascular diseases called atrial fibrillation.

### A lumped model for the cardiovascular circulation

In order to consider the cardiovascular circulation, a lumped model, which was primarily introduced by Korakianitis & Shi [14] is employed. As displayed in Figure 1, using the electric circuit analogy, the left and right hearts, and systemic and pulmonary circulations are considered. Four cardiac chambers including left atrium (LA), left ventricle (LV), right atrium (RA), and right ventricle (RV) are considered. The cardiac chambers in the model of Korakianitis & Shi [14] are modified, which will be described later on. In this study each heart chamber is defined by four components, which are representative of flow friction (*R*), inertial force (*L*), intra-cardiac compliance (*C* = 1/*E*), and flow separation coefficient (*B*). Additionally, to account for various effects of cardiac valves, the proposed model by Korakianitis & Shi [14] is adopted. To assimilate large arteries in systemic and pulmonary circulations several RCL compartments are considered. For large venous vessels given a small intravenous pressure and velocity, the inertial effects are less significant, therefore, RC components can sufficiently mimic intravenous flow attributes. Finally, the arterioles and capillaries in pulmonary and systemic circulations are incorporated through several single-element resistance components, as the resistance is the dominant flow behaviour in microcirculation. In this section the governing equations of each compartment of the lumped system will be explained in detail.

**Figure 1.**
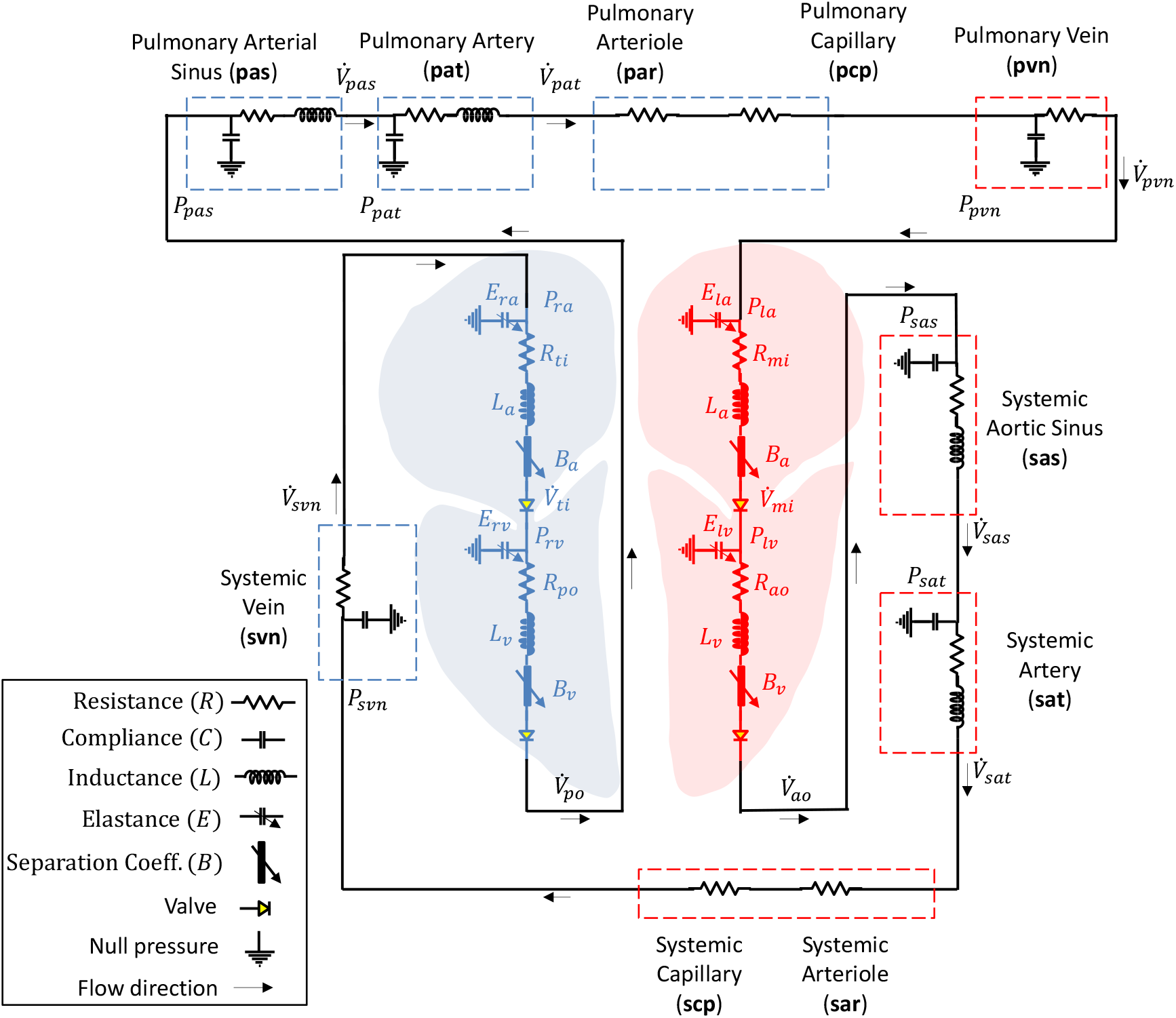
An electric circuit of a cardiovascular circulation comprising the left and the right hearts, systemic circulation, and pulmonary circulation.

The heart expansion and contraction mechanisms determine the amount of flow it can store and pass through the systemic and pulmonary circulations. Therefore, to account for the cardiac influx and efflux to each of the chambers, based on mass conservation law, the volume change of each chamber is correlated to the inflow and outflow as presented below:

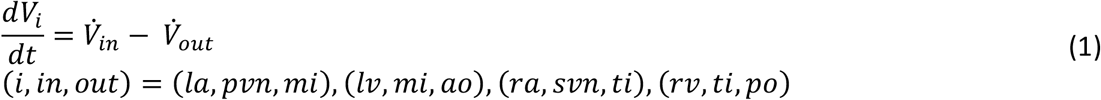

where *V* is the volume and 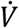 is the volumetric flow rate. Furthermore, the intra-cardiac pressure is proportional to its volume changes. The proportionality can be set to an equality by employing an appropriate elastance parameter for the heart [15]. Therefore, following equation can be written for each chamber to correlate the intracardiac pressure to its volume through appropriate elastance parameters:

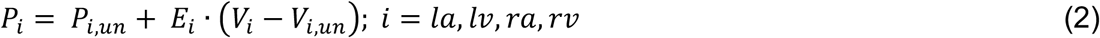

where, *P* denotes the intra-cardiac pressure and *E* is the elastance of a cardiac chamber. Moreover, subscript *un* refers to the unstressed volume and pressure of a chamber. In order to define the cardiac elastance, the following relation is proposed:

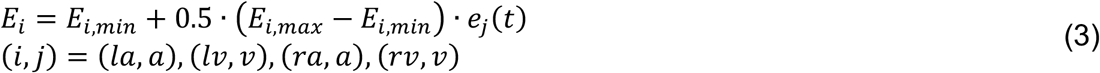

where *e*_*a*_(*t*) and *e*_*ν*_(*t*) are the normalised activation functions of the atrium and ventricle, respectively, which are used to mimic the variable heart elastance [16]. The normalised activation functions are introduced through the following equations:

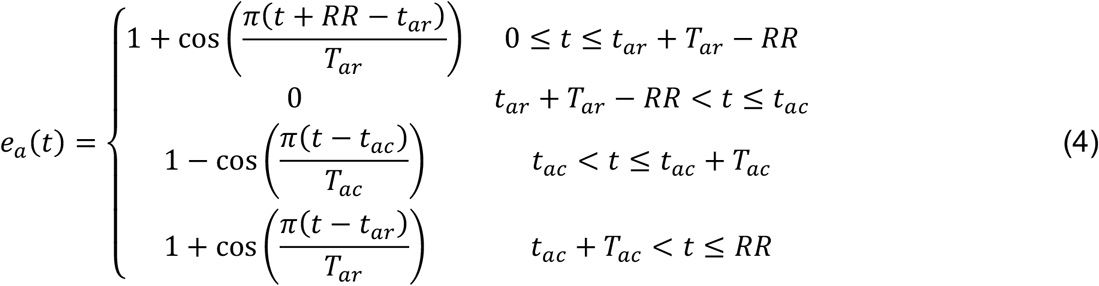

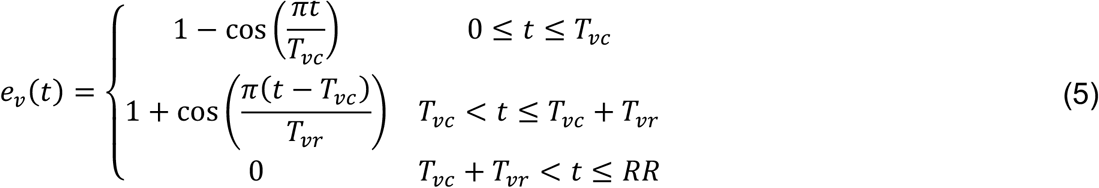

In Eqs. (4) and (5), each temporal parameter is defined as what follows:
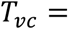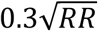 *T*_*νr*_ = 0.5 *T*_*νc*_; *T*_*ar*_ = 0.17*RR*; *T*_*ac*_ = 0.17RR; *t*_*ar*_ = 0.97*RR*; *t*_*ac*_ = 0.8*RR*.

As alluded to above, each cardiac chamber is defined by four elements. The main advantage of the selected model over its counterpart, which was suggested by Korakianitis and Shi [14] is that it eliminates the imperative iteration for finding the intracardiac flow rates. In fact, the flow across each valve is defined through a first order ordinary differential equation (ODE), which can be solved in conjunction with other ODEs. Therefore, based on the momentum equations proposed by Blanco & Feijoo [17], the following set of equations are employed for the cardiac chambers:

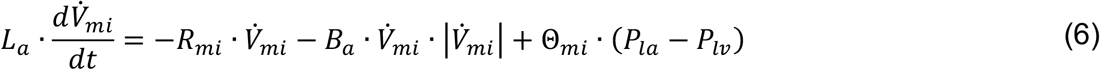

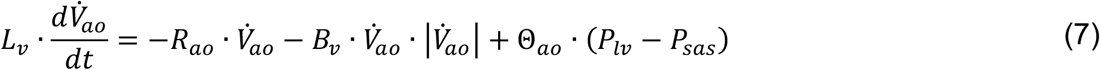

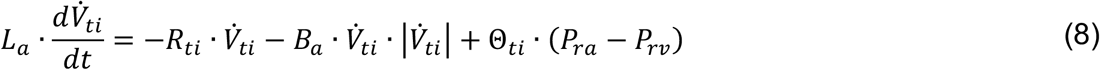

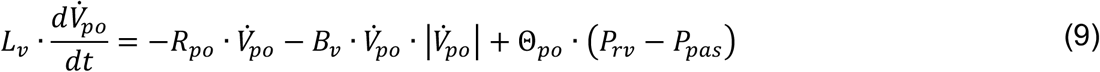

To clarify the contribution of each term in the momentum equations, they should be defined. In the left side of the equations, 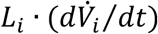 causes the pressure difference that enforcing the blood to flow, 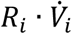 is the flow resistance due to the viscous dissipation effect, and 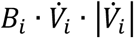 is the pressure loss due to the flow separation. Furthermore, the valve coefficient, Θ, is a function of valve opening and closing angles [17], and for each cardiac valve it can be defined as follows:

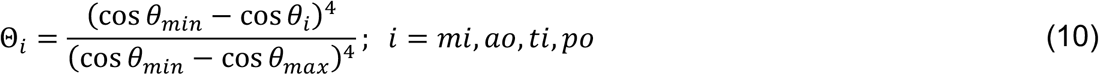

Employing a more developed valve model rather than a simple orifice with a binary motion (immediate closing and opening), a second order ODE will be resulted, which describes the valve dynamics [14]. In Eqs. (11) - (14), the first term in the right side of the equations calculates the normal pressure on the surface of a valvular leaflet. The second term considers the frictional effects due to the tissue resistance, which were assumed to be proportional to the angular motion of a valve. The third term determines the force acting on the valvular leaflet via normal velocity component of the flow, and finally, the last term considers vortex effects.

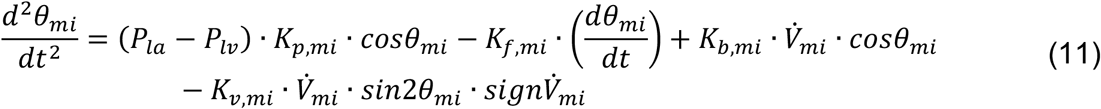

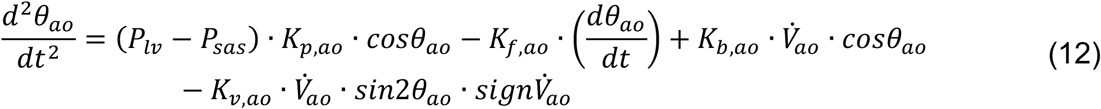

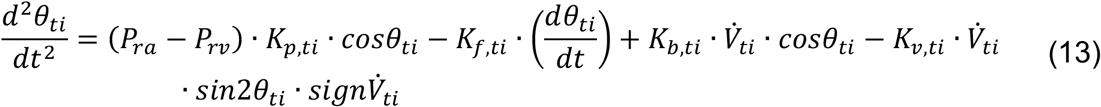

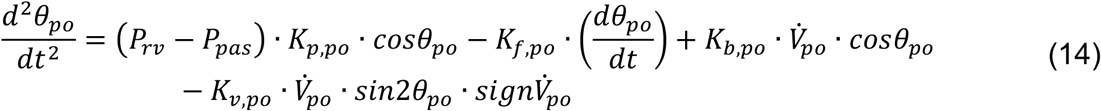

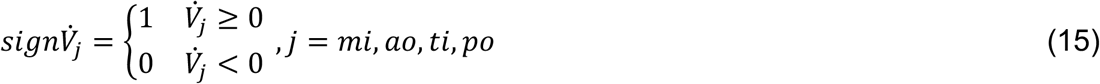

For the systemic and pulmonary loops, based on the Kirchhoff’s law for the flow rate and pressure – the former is analogous to the electrical current and the latter is analogous to the voltage, the following set of equations are obtained:

**Systemic aortic sinus**

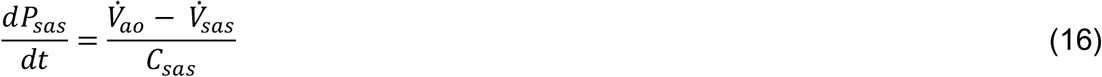

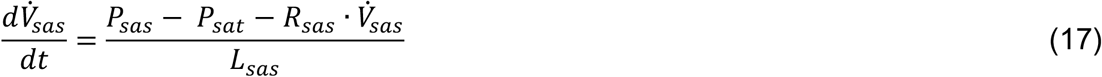
**Systemic artery**

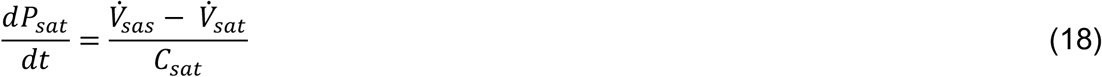

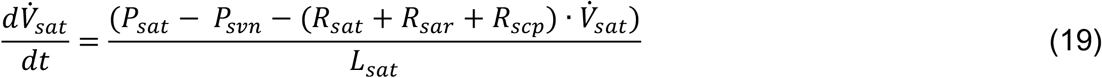
**Systemic vein**

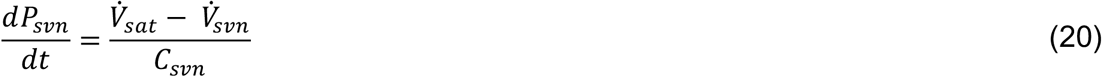

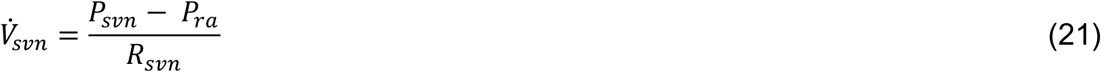
**Pulmonary arterial sinus**

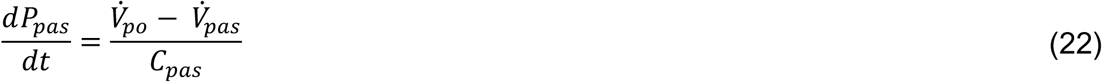

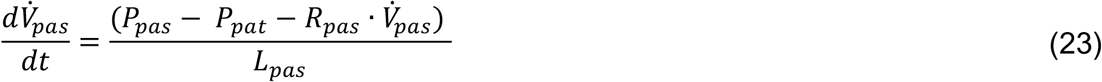
**Pulmonary artery**

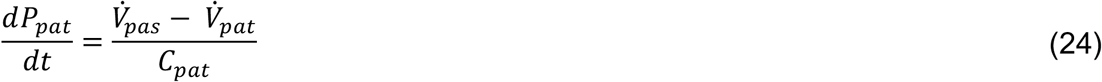

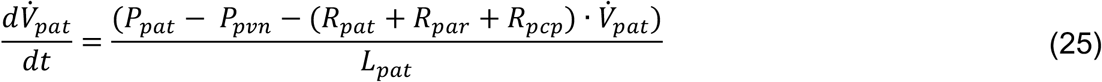
**Pulmonary vein**

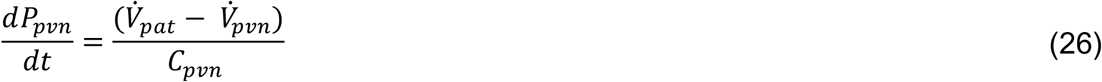

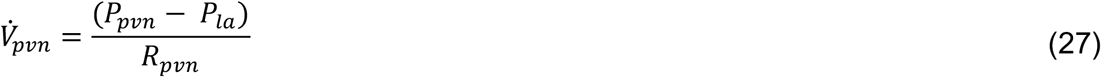

The lumped model comprises plenty of parameters, which has to be tuned appropriately to address different physiological conditions of the vascular network, particularly the heart. Table 1 presents the numeric values for a healthy candidate.

**Table 1.**
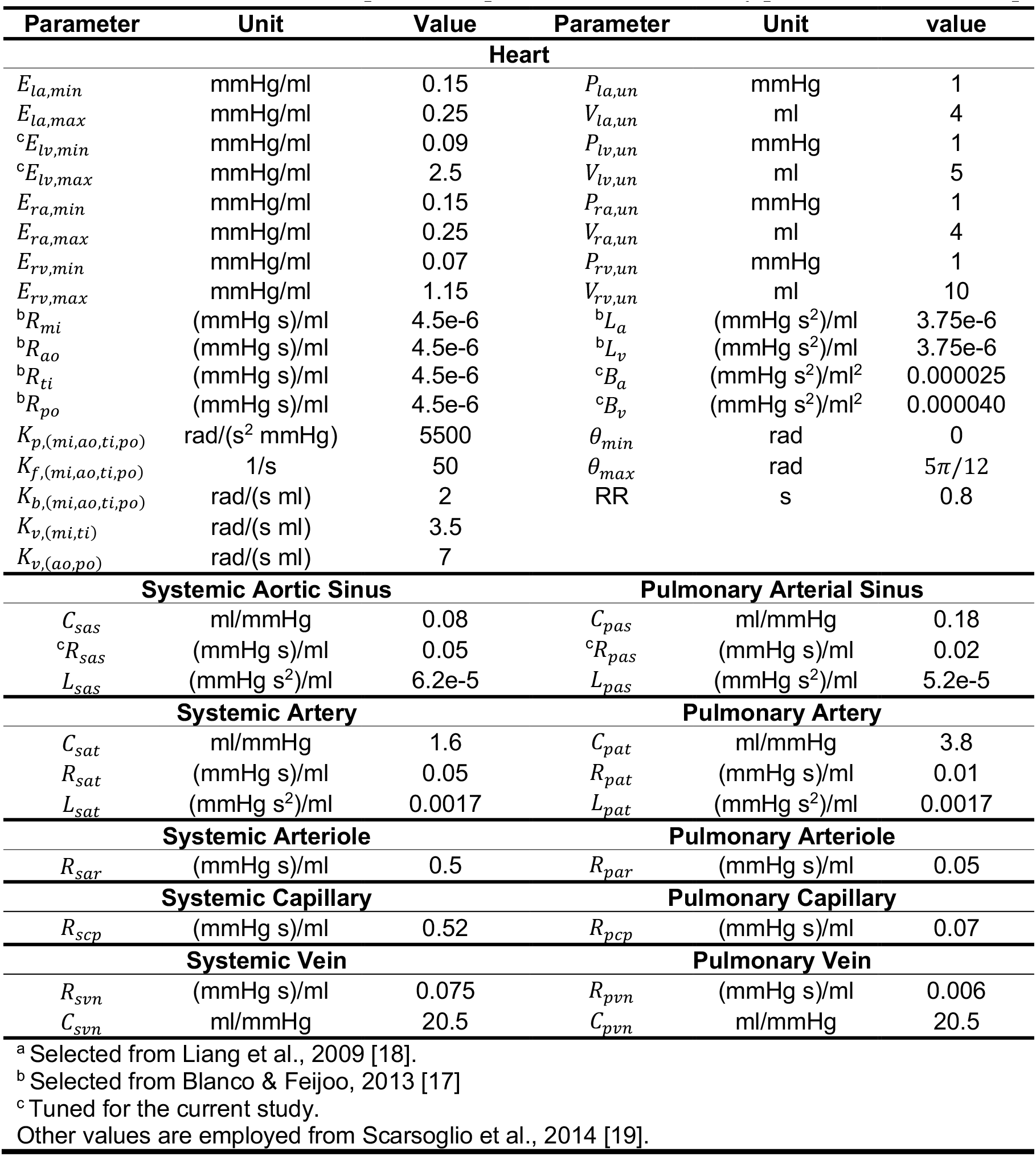
Baseline values of the lumped model parameters for a healthy person with HR = 75 bpm.

### A lumped model for the body thermoregulation

When the heat transfer and thermodynamic states of a heart is of interest, evaluating its thermal response in various physiological and environmental conditions become important. Therefore, an appropriate thermoregulation model proposes a subtle framework to measure thermal state of a heart. For that purpose, the present work is invoked the thermoregulation model proposed by Gagge et al. [3] and developed by Zhang et al. [4]. The model is known as a two-node thermoregulation model, and it consists of the core and skin regions of a body. In particular, the model can consider the temperature of skin, core, left and right hearts, artery, and vein. Although the cardiac and vascular network are not exposed to significant temperature variations comparing against the core temperature, for the sake of integrity they are considered, as separate components in the model. In response to different environmental conditions, body regulates its temperature through various mechanisms, most importantly, heart rate, vasomotion (vasodilation and vasoconstriction), shivering, and sweating. Vasomotion, shivering, and sweating work through thermal signals the body receives. In fact, each signal implies the warmness or coldness, thus, the corresponding mechanism will be activated to regulate the body temperature. To mathematically describe, these signals are defined as follows [4]:

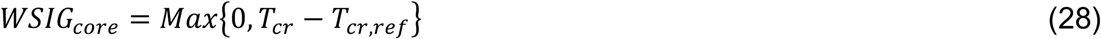

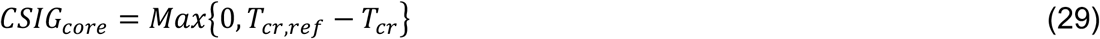

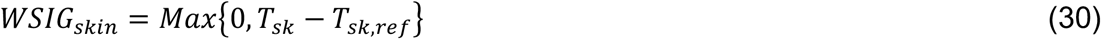

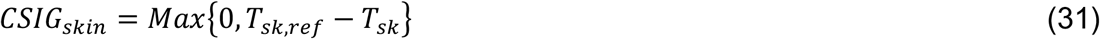

where, WSIG and CSIG are the warm and cold signals, respectively, *T*_*cr*_ (°C) is the core temperature, *T*_*sk*_ (°C) is the skin temperature. Furthermore, *T*_*cr,ref*_ (°C) and *T*_*sk,ref*_ (°C) are the reference temperatures of the core and skin, respectively. In order to incorporate different heat transfer mechanisms inside the body and through the skin, the energy balance equations for the core and skin regions are written as follows [4]:

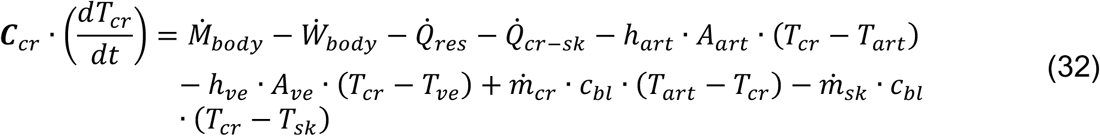

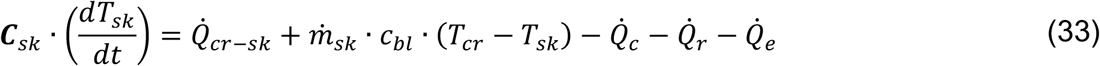

Eq. (32), the energy balance equation for the core, incorporates metabolism of the body, body work, respiration heat transfer, core to skin heat transfer, convective heat transfer between the arterial system, venal system, and core region, the heat source, and the heat sink in the core and skin regions. In Eq. (33), the energy balance equation for the skin, it comprises the heat transfer between the core and skin, skin heat source, and the skin heat transfer to the environment through the convection, radiation, and evaporation. In the following, each term and related parameters will be described meticulously.

In the energy balance equations of core and skin, ***C***_*cr*_ (W hr)/°C and ***C***_*sk*_ (W hr)/°C are the heat capacitance of core and skin, respectively, which are estimated as follows [3]:

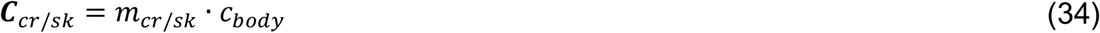

where, *C*_*body*_ (W hr °C)/kg is the specific heat of the body. The mass of core and skin are defined as 0.9584 ∙ *m*_*body*_ and 0.0416 ∙ *m*_*body*_, respectively [3]. Also, to estimate *C*_*body*_, it can be calculated through the following equation [20]:

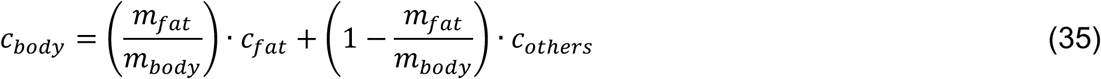

The ratio of mass of fat to the body, (*m*_*fat*_/*m*_*body*_), can be defined through the body fat (BF) percentage (%) as follows [21]:

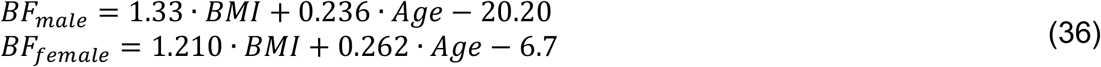

where *BMI*(kg/m^2^) is the body mass index, and *Age*(year) is the age of a person. Moreover, *h*_*ar*_ ((W K)/m^2^ and *h*_*νe*_ (W K)/m^2^ are convective heat transfer coefficients of the artery and vein, respectively, *A*_*ar*_ (m^2^) and *A*_*νe*_(m^2^) are the surface area of artery and vein, respectively, and *C*_*bl*_ (J K)/kg is the blood specific heat capacity. The rate of metabolism in the body, 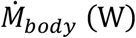 [4] is comprised of a basal metabolic rate and a shivering metabolic rate and is defined as follows:

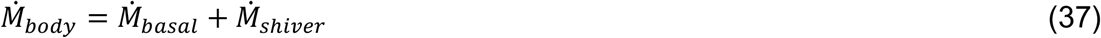

The basal metabolic rate, 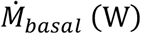, varies given the body condition and type of activity. For the body in rest, it can be estimated to be 58.2 ∙ *SA* (W) [22]. Also, the shivering metabolic rate, 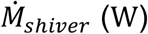, is defined as follows:

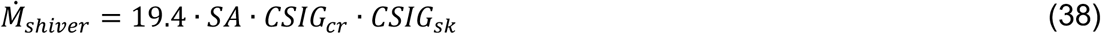

In above equations, *SA* (m^2^) is the body surface area and it can be estimated by Dubois formula [23], as presented below:

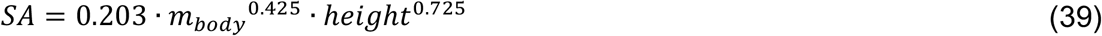

Noteworthy to mention that for the body in rest, the rate of body external work, 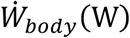, is assumed to be zero. Another heat transfer mechanism is the rate of respiration heat transfer, 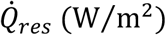 [24], which is determined as follows:

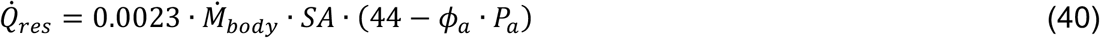

where *ϕ*_*a*_ (%) is the relative humidity, and *P*_*a*_ (mmHg) is the saturated vapour pressure of ambient air. Note that the numeric value of 44 mmHg belongs to the saturated vapour pressure at average lung temperature, which is assumed to be 35.5 °C. In general, the saturated vapour pressure can be estimated by employing the following relation [25]:

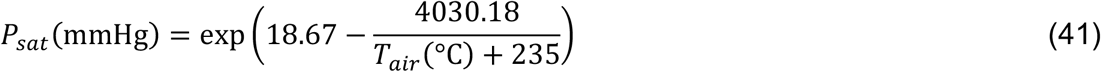

The core flow rate, 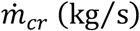, is estimated as average flow rate of systemic circulation. Moreover, skin flow rate, 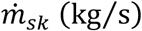, is calculated as follows [3]:

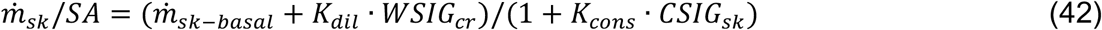

where *K*_*dil*_ kg/(m^2^ hr K) and *K*_*cons*_ (1/K) are vasodilation and vasoconstriction coefficients, respectively. Another parameter in the energy balance equations of the core and skin is the rate of heat transfer between core and skin via conduction, 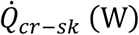, which is expressed as follows:

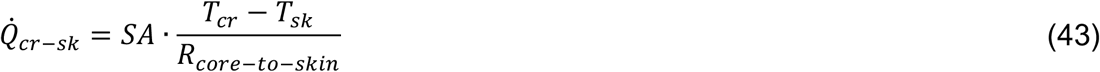

In Eq. (43), *R*_*core-to-skin*_ (m^2^ °C)/W is the thermal resistance between the core and skin, which varies in different skin blood flows and metabolic rates. In order to calculate core to skin resistance, the following relation [20] is incorporated:

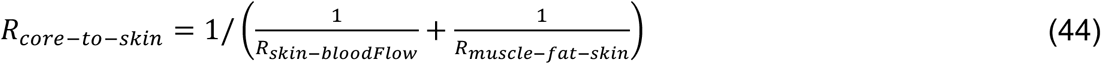

where *R*_*skin-bloodFlow*_ (m^2^ °C)/W and *R*_*muscle-fat-skin*_ (m^2^ °C)/W are defined as follows:

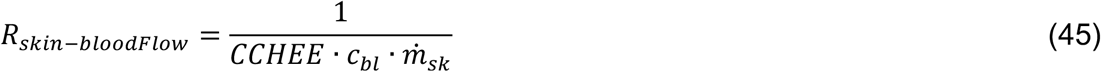

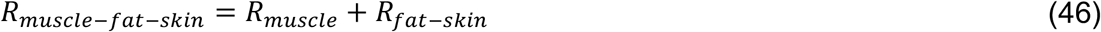

Where *CCHEE* is the counter-current heat exchanger efficiency. Furthermore, muscle-fat-skin resistance is split into *R*_*muscle*_ (m^2^ °C)/W and *R*_*fat-skin*_ (m*2* °C)/W as follows:

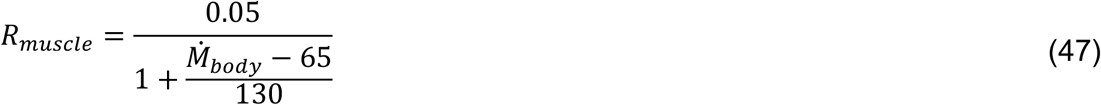

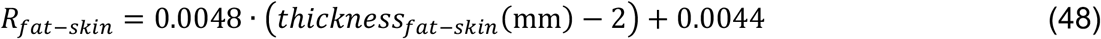

In the skin energy balance equation, 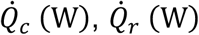, and 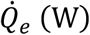 are the skin heat transfer mechanisms to the environment through the convection, radiation, and evaporation, respectively. The rate of convection and radiation heat transfers are determined as follows:

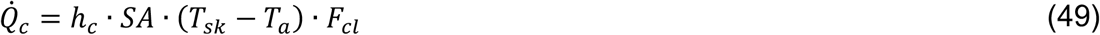

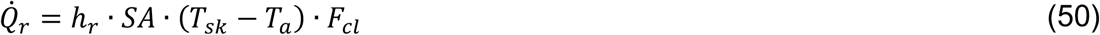

where, *h*_*c*_ W/(m^2^K) and *h*_*r*_ W/(m^2^K) are the convective and radiative heat transfer coefficients, respectively. In the present study, *h*_*r*_ is assumed to be constant, which is equal to 4.65 W/(m^2^K) [4] and *h*_*c*_ is expressed as follows [26]:

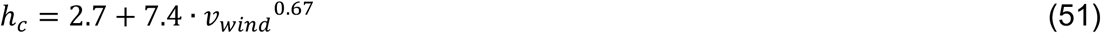

where, *ν*_*wind*_ (m/s) is the wind velocity. Also, in Eqs. (49) and (50), *F*_*cl*_ is Burton’s efficiency factor [27] for clothing, which is estimated as follows:

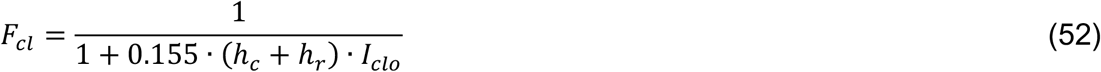

where *I*_*clo*_ (clo) is the thermal resistance coefficient. In order to clarify the associated unit of thermal resistance coefficient, 1 clo is equal to 0.155 (m^2^ °C)/W.

Another heat transfer mechanism in the skin energy balance equation is the evaporative heat loss. The evaporation from the skin surface occurs through two heat loss mechanisms [3], namely, the heat loss due to the sweating, 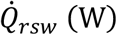, and the heat loss due to the skin diffusion, 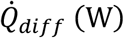. Thus, the cumulative value forms the overall rate of evaporation heat loss as defined below:

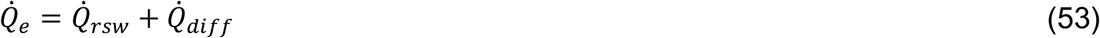

Therefore, in the above equation, rate of sweating heat transfer [3] is defined as follows:

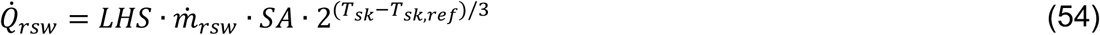

where *LHS* (W hr)/gr is the latent heat of sweat and is taken as a constant parameter, and 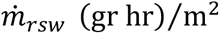 is the sweating flow rate, which is defined as follows:

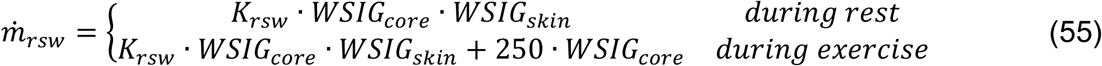

where, *K*_*rsw*_ gr/(m^2^ hr K) is the sweating rate coefficient. In order to determine the rate of diffusion heat loss [3], the following relation is employed:

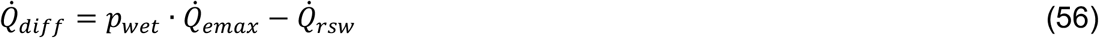

where, *P*_*wet*_ is the skin wetness, which is calculated as follows:

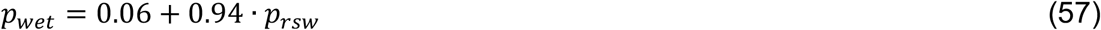

In the skin wetness equation, *p*_*rsw*_ is introduced as the ratio of sweating heat loss 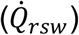 to the maximal evaporative loss 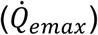, therefore, it varies between zero and unity. Furthermore, 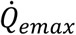 is defined as follows:

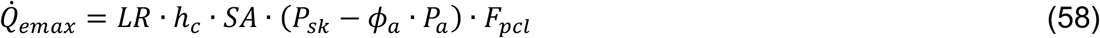

where *LR* (°C/mmHg) is the Lewis relation, *P*_*sk*_ (mmHg) is the saturated vapour pressure at the mean skin temperature (*T*_*sk*_), which can be estimated through Eq. (41). Moreover, *F*_*pcl*_ is called the permeation efficiency factor, which defines the efficiency of water vapour infiltration through the cloth to the environment as it evaporates from the skin surface. The permeation efficiency factor is estimated as follows [27]:

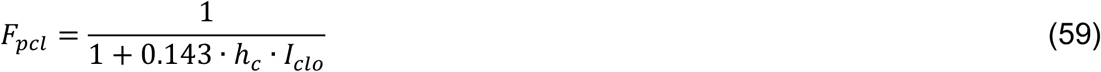

Despite the marginal difference between the core body temperature and the heart, the energy balance relation for the vascular circulations, the left heart, and the right heart are considered as follows [4]:

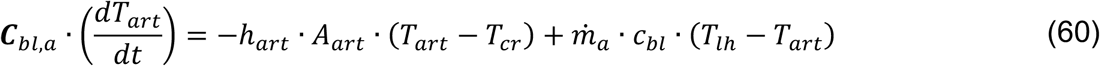

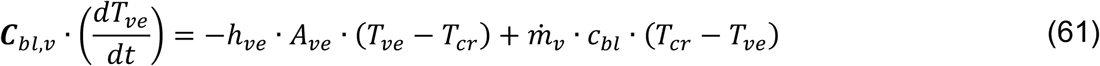

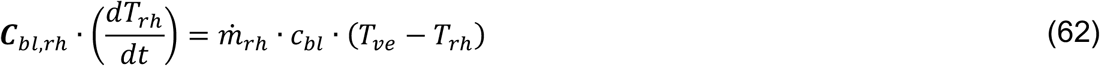

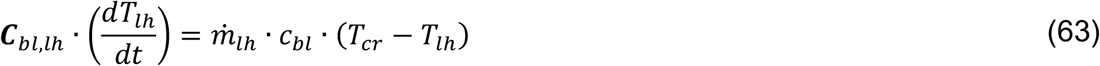

In the above-mentioned energy balance equations, *T*_*art*_(°C), *T*_*νe*_(°C), *T*_*rh*_(°C), and *T*_*lh*_ (°C) are the temperatures of systemic artery, systemic vein, right heart, and left heart, respectively. Additionally, *A*_*art*_(m^2^) and *A*_*νe*_ (m^2^) are the surface areas of systemic artery and systemic vein, respectively. Finally, *C*_*bl,a*_ (J/K) is the thermal capacitance of arteries, *C*_*bl,ν*_ (J/K) denotes the thermal capacitance of veins, *C*_*bl,rh*_ (J/K) is the thermal capacitance of right heart, and *C*_*bl,lh*_ (J/K) defines the thermal capacitance of left heart. All thermal capacitances can be calculated through the following equations:

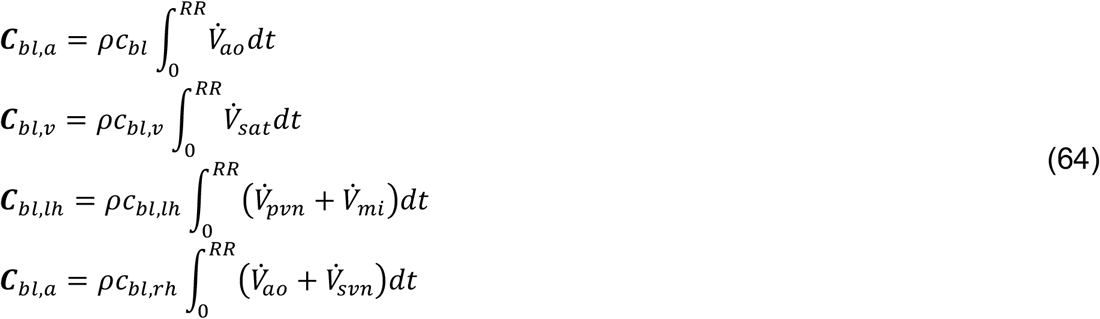

Note that, the mass flow rate is simply obtained as follows:

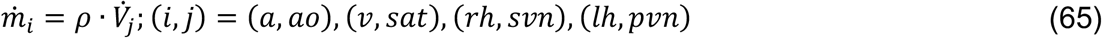

Table 2 presents the employed parameters, with their corresponding units and numeric values in a constant environment conditions and the body at rest.

**Table 2.** Parameters of the body thermoregulation model.

| Parameter definition | Symbol | Unit | Value |
| --- | --- | --- | --- |
| Core reference temperature | $T_{cr,ref}$ | °C | 36.8 |
| Skin reference temperature | $T_{sk,ref}$ | °C | 34.1 |
| Total mass | $m_{body}$ | kg | 81.7 |
| Height | $height$ | m | 1.80 |
| Age | $age$ | year | 24 |
| Specific heat capacity of fat | $c_{fat}$ | J/(kg K) | 2510 |
| Specific heat capacity of body without fat | $c_{other}$ | J/(kg K) | 3650 |
| Ambient temperature | $T_a$ | °C | 25 |
| Relative humidity | $\phi_a$ | % | 47 |
| Basal Metabolism | $\dot{M}_b$ | w | $58.2 \cdot SA$ |
| Surface area of systemic artery | $A_{art}$ | m<sup>2</sup> | 0.12 |
| Surface area of systemic vein | $A_{ve}$ | m<sup>2</sup> | 0.2 |
| Convection heat transfer coefficient of artery | $h_{art}$ | W/(m<sup>2</sup>K) | 1800 |
| Convection heat transfer coefficient of vein | $h_{ve}$ | W/(m<sup>2</sup>K) | 1080 |
| Specific heat capacity of blood | $c_{bl}$ | J/(kg K) | 3640 |
| Blood density | $\rho_{bl}$ | kg/m<sup>3</sup> | 1060 |
| Basal skin blood flow | $\dot{m}_{sk-basal}$ | kg/(m<sup>2</sup> hr) | 6.3 |
| Vasodilation coefficient | $K_{dil}$ | kg/(m<sup>2</sup> hr K) | 75 |
| Vasoconstriction coefficient | $K_{cons}$ | 1/K | 0.5 |
| Radiative heat transfer coefficient | $h_r$ | W/(m<sup>2</sup> K) | 4.65 |
| Wind velocity | $v_{wind}$ | m/s | 0.2 |
| Thermal resistance of clothing<br>(insulation of clothing) | $I_{clo}$ | clo | 0.1 |
| Latent heat of sweat | $LHS$ | (W hr)/gr | 0.7 |
| Sweating rate coefficient | $K_{rsw}$ | gr/(m<sup>2</sup> hr K) | 100 |
| Lewis relation | $LR$ | °C/mmHg | 2.2 |
| Thickness fat-skin | $thickness_{fat-skin}$ | mm | 15.72 |

### Numerical method, solving the governing ODEs

The coupled lumped model comprises two sets of ODEs, which expresses the cardiovascular flow and body thermal condition. The flow lumped model results 22 ODEs for finding flow rate, pressure, and angular position of the valves. Also, the energy balance equations of the body thermoregulation led to 6 ODEs to estimate the temperatures.

The set of equations is solved chronologically as shown in Figure 2. In fact, initially the flow lumped model ODEs are solved, then the resulted outputs will be passed through the thermoregulation model to find the temperatures. Furthermore, there was employed an auxiliary block, which takes in flow data for further necessary calculations. The output of the coupled flow-thermoregulation lumped model will be employed for further analysis of cardiac function or as the boundary condition of flow as it leaves the LV. The code was developed in MATLAB R2020a, and a MATLAB ODE solver, ode15s, was invoked. Ode15s is a robust variable-step variable-order method based on the numerical discretisation formulas, which can effectively resolve stiff equations.

**Figure 2.**
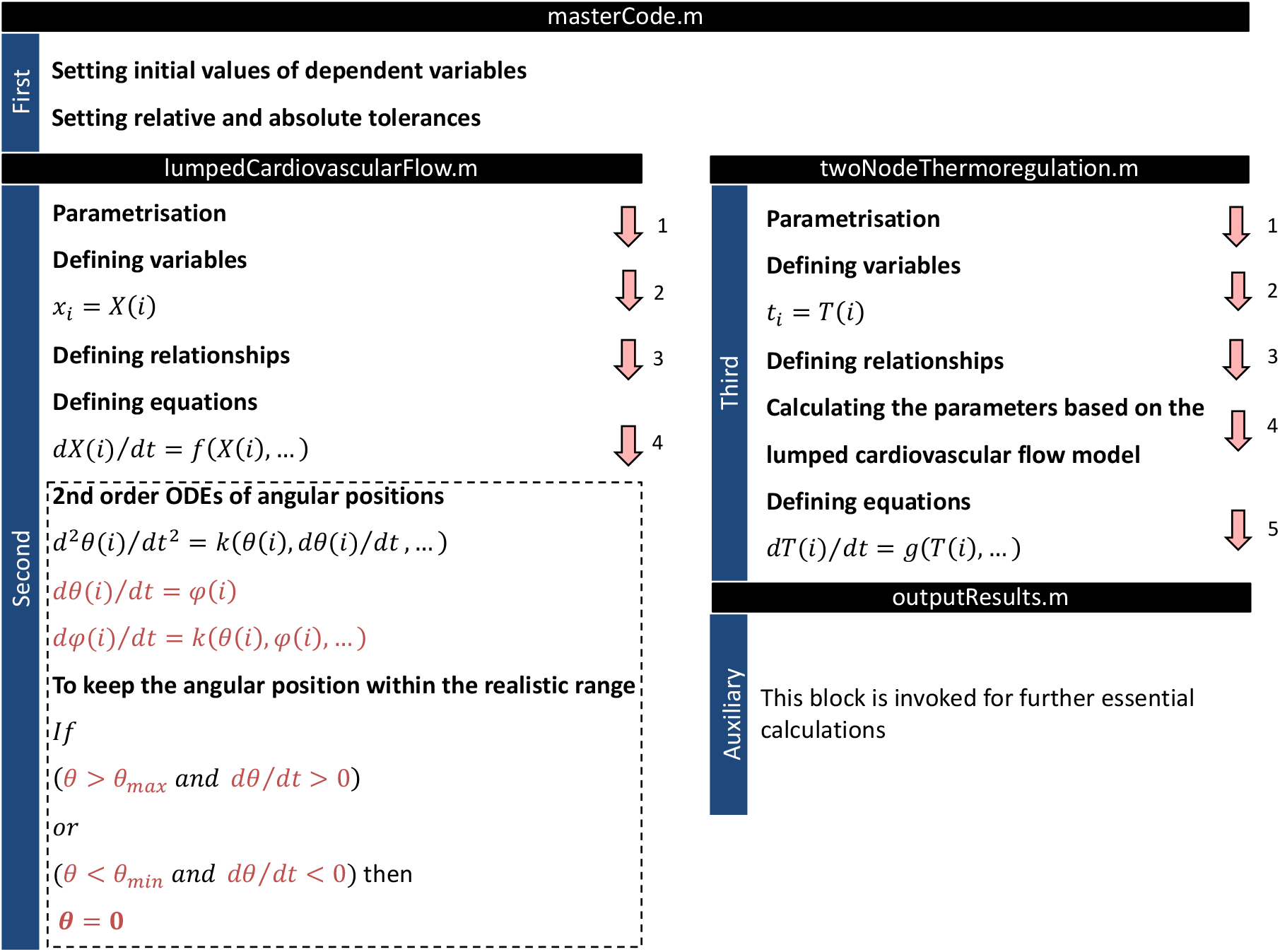
Solution algorithm for governing equations of lumped flow and thermoregulation models.

### Initial values of the governing equations

**Table 3.**
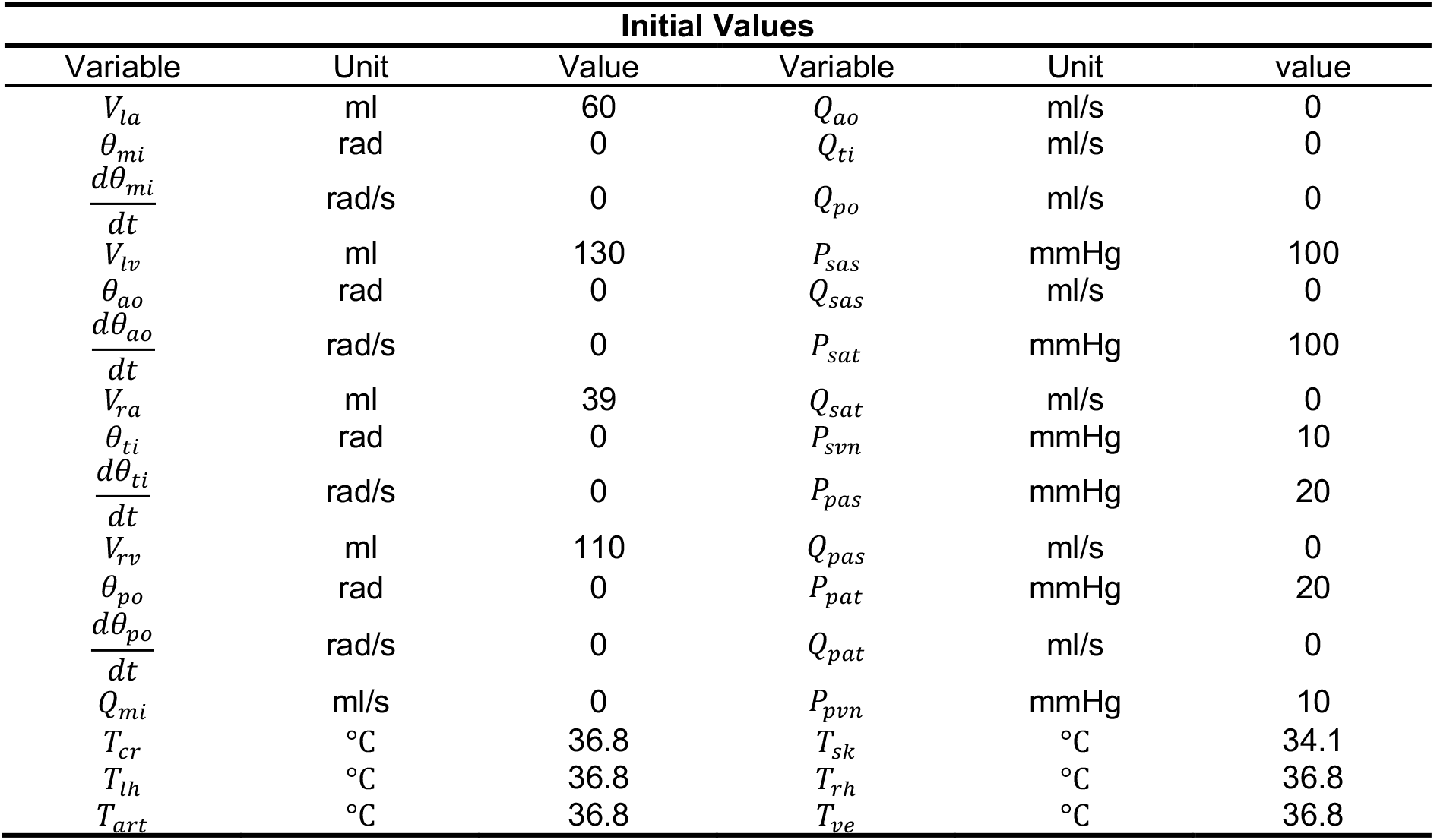
Initial values for solving governing equations of the lumped flow and thermoregulation models.

### Validation of the Models

In this section to evaluate the developed pipeline, serval comparisons are made against experimental data and in-vivo measurements. Figure 3(a) and (b) show intra-cardiac and aortic pressures. Figure 3(c) and (d) demonstrate the flow rates across the four cardiac valves. Furthermore, Figure 3(e) illustrates the volume changes of heart chambers, and finally in Figure 3(f), LV pressure-volume relation is plotted for different preload (*R*_*mitral*_) conditions.

**Figure 3.**
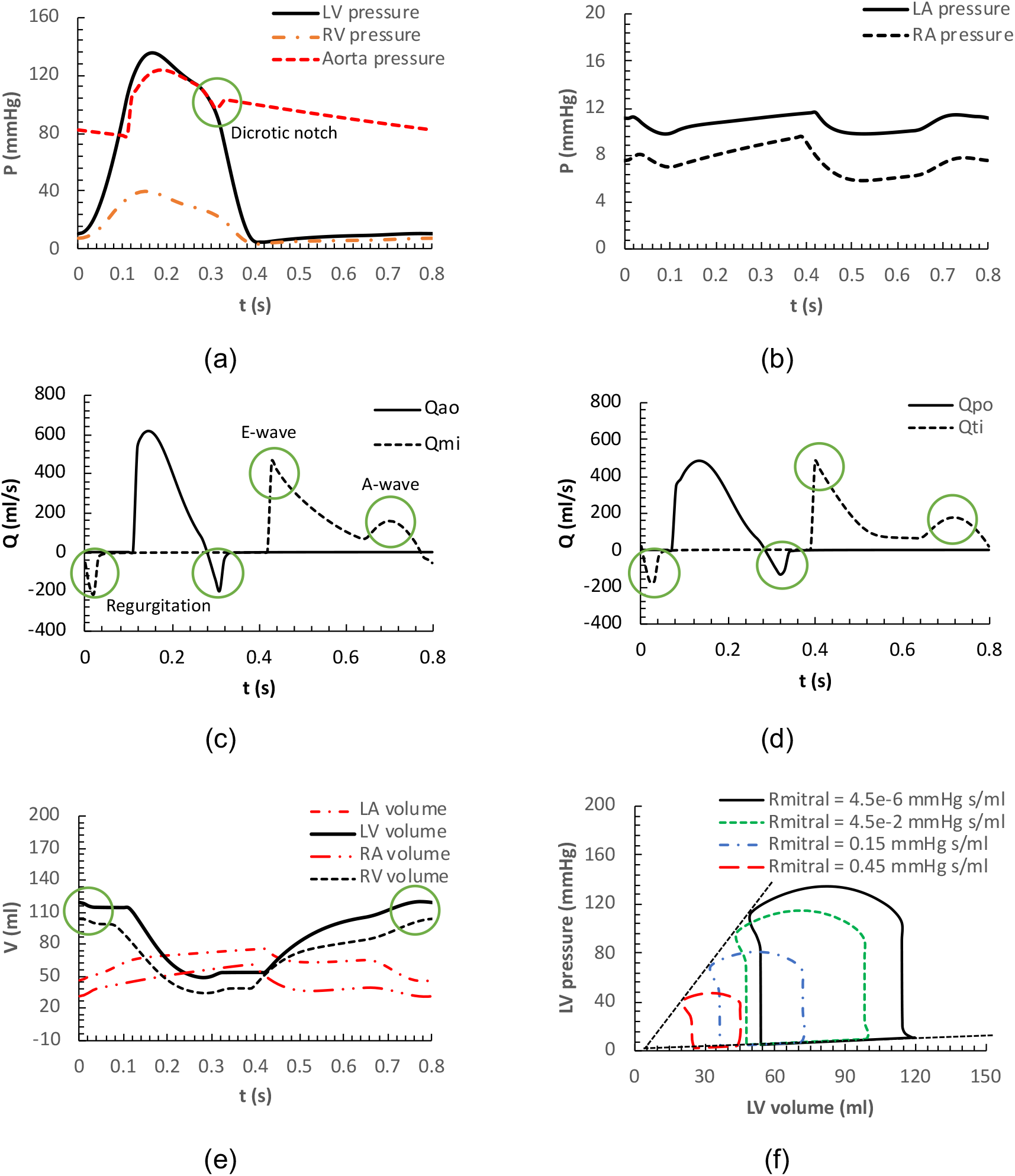
(a) pressure of the left ventricle (LV), right ventricle (RV), and aorta in a cardiac cycle, (b) pressure of the left atrium (LA) and right atrium (RA) in a cardiac cycle, (c) flow rates across aortic and mitral valves, (d) flow rates across pulmonary and tricuspid valves, (e) volume changes of the LA, LV, RA, and RV in a cardiac cycle, and (f) LV pressure-volume relation for different preload (Rmitral) conditions.

As shown in Figure 3(a) and (b), the left heart pressure is higher than the right heart pressure, and the ventricular pressure is higher than the atrial pressure. The results emphasise that for a normal heart function, the LA varies between 9 and 12 mmHg and the LV changes between 5 and 134 mmHg. While the values for the RA (5-9 mmHg) and RV (3-40 mmHg) are less than the left heart. Also, about the aortic pressure waveform in Figure 3(a), it shows that the model can predict the aortic notch, when the aortic valve opens. In Figure 3(c) and (d) the flow rates across the aortic, mitral, pulmonary, and tricuspid valves are displayed. The model can successfully predict important flow features, mainly, the flow regurgitation during the valvular closure, and E-wave and A-wave peaks across the mitral and tricuspid valves. Note that E-wave and A-wave indicate the passive and active contractions of normal atria, respectively. Therefore, the obtained data are in a physiological range reported for normal people and they are in a good agreement with other numerical studies [14], [28].

Figure 3(e) displays the volume changes of different chambers. The current model suggests that the stroke volumes for the LA and LV are about 30 ml and 70 ml, respectively, which are well located in a range reported by Gutman et al. [29] for a healthy cohort (24 ± 7.6 for LA and 65 ± 14.3 for LV). Furthermore, the cardiac output is about 5.2 l/min which is in a normal range reported in textbooks of physiology [30]. End systolic pressure volume relation (ESPVR) as one of the well-known heart parameter [31] is plotted in Figure 3(f) for different preload conditions. The pressure-volume diagram of the LV displays a good agreement with the real physiological conditions. In fact, the loci of end systolic pressure volume result a straight line, which intercepts the abscissa at the value of LV unstressed volume.

In addition to the flow lumped model, the thermoregulation model was also validated against a set experimental data. Figure 4(a) and (b) compare the result of the current study against the experimental data obtained by Takada et al. [32]. The comparison is made for two schedules and subject D of the study by Takada et al. [32]. In schedule I, everyone underwent various thermal conditions in five different timespans, while in schedule II, the cases were tested in three different thermal conditions. To improve the comparison, the thermal signal reference temperatures, vasomotion parameters, and basal skin blood flow were tuned to be consistent with thermal characteristics of people in the experiment. The results of the employed thermoregulation model show a good agreement with the experimental data.

**Figure 4.**
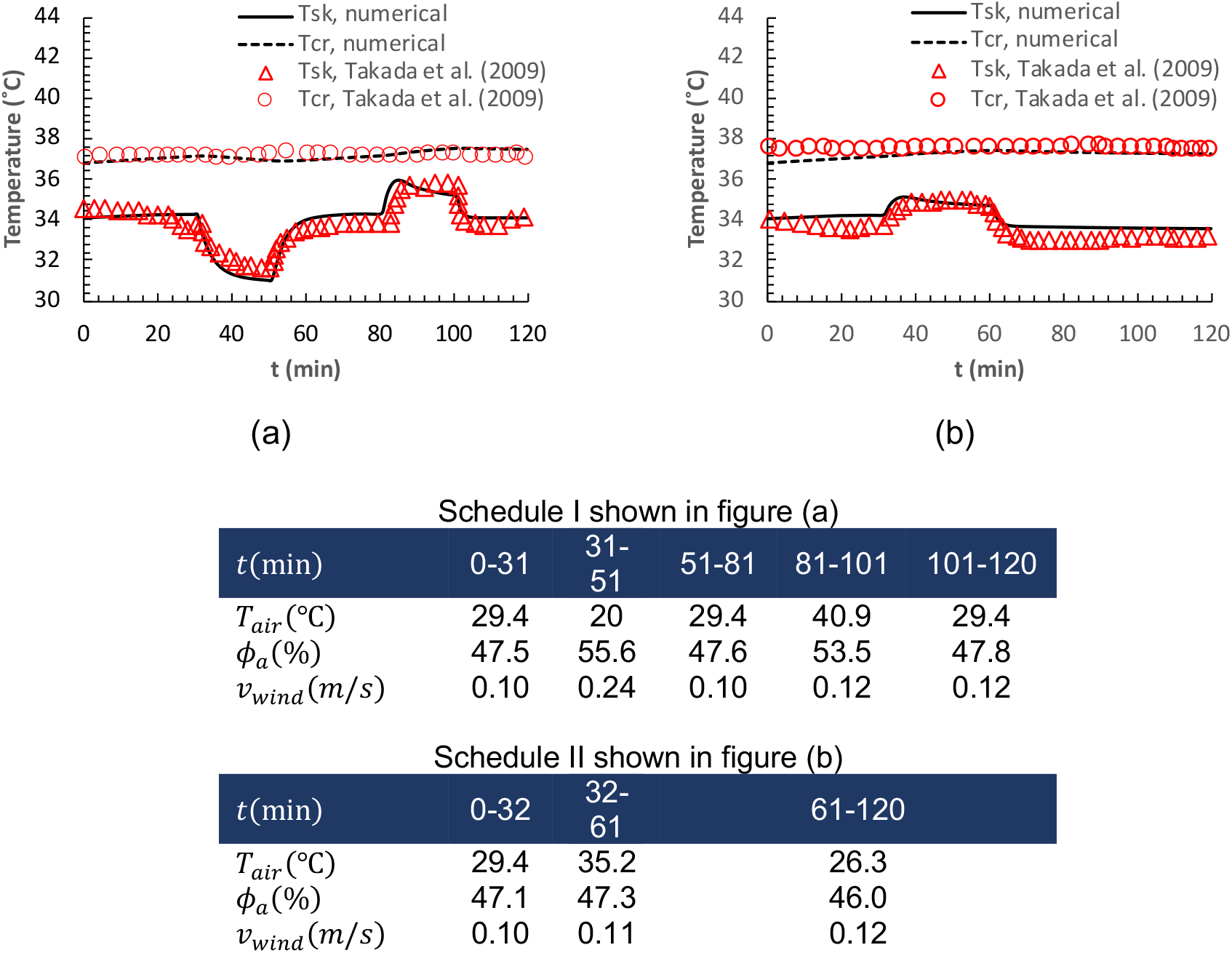
Skin and core temperatures, comparing the numerical results against the experimental data of subject D in the study by Takada et al. [32], (a) schedule I, and (b) schedule II.

## Conclusion

In this methodology paper, a workflow was developed based on previous lumped models of flow and body’s thermoregulation. The develop pipeline considers the heart, systemic, and pulmonary circulations of blood flow, while it accounts for the thermal response of the body. The coupled pipeline provides crucial information about flow rate, pressure, and temperature of cardiovascular system. The pipeline was successfully validated against several clinical and experimental dataset to evaluate its credential. The outcome of this model can be interpreted for variety of physiological conditions and provide essential information for further analysis. The workflow is mainly developed to explore effects of atrial fibrillation on cardiac function. Furthermore, it provides the necessary input data for entropy and exergy analysis of the cardiac system, which consequently leads to a new index for the heart ageing.

